# Broadening sarbecovirus neutralization with bispecific antibodies combining distinct conserved targets on the receptor binding domain

**DOI:** 10.1101/2024.05.09.593284

**Authors:** Denise Guerra, Laura Radić, Mitch Brinkkemper, Meliawati Poniman, Lara van der Maas, Jonathan L. Torres, Andrew B. Ward, Kwinten Sliepen, Janke Schinkel, Rogier W. Sanders, Marit J. van Gils, Tim Beaumont

## Abstract

Monoclonal neutralizing antibodies (mAbs) are considered an important prophylactic against SARS-CoV-2 infection in at-risk populations and a strategy to counteract future sarbecovirus-induced disease. However, most mAbs isolated so far neutralize only a few sarbecovirus strains. Therefore, there is a growing interest in bispecific antibodies (bsAbs) which can simultaneously target different spike epitopes and thereby increase neutralizing breadth and prevent viral escape. Here, we generate and characterize a panel of 30 novel broadly reactive bsAbs using an efficient controlled Fab-arm exchange protocol. We specifically combine some of the broadest mAbs described so far, which target conserved epitopes on the receptor binding domain (RBD). Several bsAbs show superior cross-binding and neutralization compared to the parental mAbs against sarbecoviruses from diverse clades, including recent SARS-CoV-2 variants. BsAbs which include mAb COVA2-02 are among the most potent and broad combinations. As a result, we study the unknown epitope of COVA2-02 and show that this mAb targets a distinct conserved region at the base of the RBD, which could be of interest when designing next-generation bsAb constructs to contribute to a better pandemic preparedness.

## Introduction

Sarbecoviruses are a subgenus of the Coronaviridae family with a high potential to cause spillover events from animal reservoirs to humans, resulting in serious respiratory diseases and leading to global health emergencies ^1^. Recently, they have gained worldwide attention as Severe Acute Respiratory Syndrome Coronavirus 2 (SARS-CoV-2), a sarbecovirus from the Betacoronavirus genus, caused the global Coronavirus Disease 2019 (COVID-19) pandemic ^2,3^.

Since the beginning of the pandemic, enormous efforts have been put into the isolation, characterization and large-scale production of monoclonal antibodies (mAbs) for therapeutic and prophylactic purposes. Multiple mAbs have been isolated from infected individuals during the first wave of the pandemic, and some of them have initially shown efficacy against the first variants of SARS-CoV-2 ^4,5^. However, with the more recent emergence of highly resistant viral strains, such as the Omicron sub-lineages belonging to a different antigenic cluster ^6^, most of these mAbs have shown reduced or completely abolished efficacy ^7–9^.

What the most successful antibody therapeutics have in common is their breadth in targeting multiple SARS-CoV-2 variants, which often extends to other members of the sarbecovirus subgenus. These include SARS-CoV, which caused an epidemic in 2003 ^1^, SHC014, Pangolin GX 2017, Rf1 and Khosta-2, found in animal hosts including bats and pangolins ^10^. Besides their valuable role in protecting or treating immunocompromised and other vulnerable populations who can not receive protective vaccines, mAbs with broad reactivity to existing sarbecoviruses can be considered a proxy for quickly combating future outbreaks and could thus be crucial tools for pandemic preparedness.

The cross-reactivity of these mAbs can be explained by the conserved epitopes they target on the trimeric SARS-CoV-2 spike (S) glycoprotein. These epitopes are the membrane proximal S2 subunit, with the fusion peptide and the heptad repeats 1 and 2, or regions at the base of the receptor binding domain (RBD), distant from the more exposed and mutation-prone angiotensin converting enzyme 2 (ACE2) binding site ^11–15^. Despite their broad reactivity, S2-binding mAbs do not often show strong neutralization ^16,17^. In contrast, mAbs targeting the lower part of the RBD have been reported to be more efficacious at neutralizing SARS-CoV-2 than S2-binding mAbs, although usually less potent than mAbs targeting upper RBD regions ^12,15^. Since broad RBD-targeting mAbs do not directly interfere with ACE2 binding, their mechanisms of action often include destabilization of the S protein, intra- and inter-S crosslinking (i.e. binding to two epitopes on one or two S simultaneously) and driving the RBDs into conformational changes that sterically hinder receptor binding ^18,19^.

Due to viral resistance to natural and vaccine-induced immunity, and to many of the previously described mAbs ^7–9^, additional strategies to combat SARS-CoV-2 have been investigated. One of these approaches is combining the properties of multiple mAbs into bivalent and multivalent antibody-like constructs, such as bispecific antibodies (bsAbs). Different SARS-CoV-2 targeting bsAbs have been described, which show improved neutralization activity compared to the parental mAbs ^18,20,21^. The so far described IgG-like bsAbs are of highly diverse formats, such as the CrossMAb constructs ^22–26^, DVD-Ig ^24,27^, IgG-(scFv)2 ^21,24,28,29^, Tandem scFv-Fc ^28,30,31^, and VH/Fab IgGs ^32,33^, many of which require significant engineering and quality control processes to be efficiently produced. Additionally, most of these bsAbs usually target at least one exposed and non-conserved epitope, which can make them susceptible to viral escape. Therefore, considering the continued SARS-CoV-2 circulation and emergence of new variants, as well as the potential for zoonotic spillover of other sarbecoviruses to the human population, a need for novel broad (bs)Ab therapeutics remains.

Here, we used controlled Fab-arm exchange (cFAE) ^34^ to generate a panel of 30 novel IgG1-like bsAbs. In this method, single point mutations are introduced in the Fc regions of each parental mAb. After individual purification of the mAbs and co-incubation with a reducing agent at 31°C for 5 h, the Fab arms get spontaneously exchanged, forming bsAbs with a high efficacy (>90%). To create bsAbs, we selected some of the broadest RBD-targeting mAbs described so far. MAbs targeting the RBD are categorized into different classes based on their epitope ^15,35^. We specifically focused on class 3 and 4 mAbs as they are directed towards the more conserved base of the RBD ^11–15^. Additionally, we included potent candidates isolated from participants of the COSCA cohort in Amsterdam which showed neutralizing breadth ^4,20^. Some of our bsAb candidates demonstrate enhanced breadth and improved potency in neutralization assays against a panel of SARS-CoV-2 variants and other sarbecoviruses. This work substantiates the hypothesis that by simultaneous targeting of non-overlapping conserved RBD epitopes with bsAbs, it is possible to achieve increased neutralizing breadth. BsAbs could thus be a promising countermeasure for future mutational variants and potential sarbecovirus outbreaks.

## Materials and Methods

### Cell lines

Human embryonic kidney (HEK)293T (ATCC CRL-11268) and FreeStyle 293F (Life Technologies) cell lines were used for recombinant protein productions. Briefly, HEK293T cells were cultured in DMEM (Sigma-Aldrich) medium supplemented with 10% fetal calf serum (FCS) and 1% penicillin-streptomycin at 37°C with 5% CO_2_, and maintained at a confluency of 40-80%, while FreeStyle 293F cells were cultured in FreeStyle 293 expression medium (Thermo Scientific) at 37°C with 8% CO_2_. For pseudovirus neutralization assays, HEK293T expressing the human ACE2 receptor were employed. HEK293T/ACE2 cells were cultured in DMEM + 10% fetal bovine serum (FBS) and 1% penicillin-streptomycin at 37°C with 5% CO_2_.

### Constructs design

Soluble, prefusion SARS-CoV-2 and sarbecovirus S proteins were generated as described previously ^4^. In short, a gene encoding residues 1-1138 with proline substitutions at positions 986 and 987 and a GGGG substitution at positions 682-685 was ordered as gBlock gene fragment (Integrated DNA Technologies) and cloned into a pPPI4 plasmid backbone containing a T4 trimerization domain with Gibson Assembly (ThermoFisher), followed by a hexahistidine (his) tag. Prefusion Omicron BA.2 and BA.4/5 S proteins were kindly provided by Dirk Eggink of the National Institute for Public Health and the Environment (RIVM), Bilthoven, the Netherlands. Compared to the WT S protein, the following mutations were present in the variant proteins: Omicron BA.2 (T19I, L24S, Δ25/27, G142D, V213G, G339D, S371F, S373P, S375F, T376A, D405N, R408S, K417N, N440K, S447N, T478K, E484A, Q493R, Q498R, N501Y, Y505H, D614G, H655Y, N679K, P681H, N764K, D796Y, Q954H, N969K), Omicron BA.4/5 (T19I, L24S, Δ25-27, Δ69-70, G142D, V213G, G339D, S371F, S373P, S375F, T376A, D405N, R408S, K417N, N440K, L452R, S477N, T478K, E484A, F486V, Q498R, N501Y, Y505H, D614G, H655Y, N679K, P681H, N764K, D796Y, Q954H, N969K). COVA mAbs used in the study (COVA2-02, COVA1-16, COVA1-18, COVA2-15, COVA1-22, COVA309-22, COVA309-35 and COVA309-38) were isolated from participants included in the COSCA study (NL73281.018.20) ^4^. Briefly, the variable V(D)J-regions of the heavy and light chain (HC and LC) of the mAbs were cloned into corresponding expression vectors containing the constant regions of the human IgG1, as previously described ^4^. The variable HC and LC sequences of mAb S309, Ly-CoV1404, CV38-142, CR3022 and 10-40 were ordered as gene fragments and cloned into the same corresponding expression vectors for HC and LC, respectively. To generate Fab fragments, mAbs were incubated at 37□ for 5 h with 100 µL papain immobilized on agarose resin (ThermoFisher Scientific) in PBS, 10 mM EDTA, 20 mM cysteine at pH 7.4. Subsequently, the Fc domain and non-digested mAbs were removed from the flow-through by a 2 h incubation at room temperature with 200 µL of protein A resin (ThermoFisher Scientific) per mg of mAb. Finally, the mix was centrifuged in a Costar Spin-X filter centrifuge tube (Corning) and flow-through containing the Fabs was buffer exchanged to TBS using Vivaspin filters with a 10 kDa molecular weight cutoff (GE Healthcare).

### Production of viral proteins and mAbs

After constructs verification by Sanger sequencing, all soluble S proteins were produced in HEK293F suspension cells (ThermoFisher), followed by affinity purification using Ni-NTA agarose beads and size exclusion chromatography (SEC), as described previously ^4^. To produce mAbs, co-transfection of the HC and LC plasmids in HEK293F cells was performed in a 1:1 ratio and harvested after 5 days. The filtered supernatants were run over 10 ml protein G columns (Pierce). After elution, the purified mAbs were buffer exchanged to PBS using 100kDa Vivaspin6 columns. NanoDrop One (Thermofisher) was used to measure the IgG concentration. All mAbs were then diluted with PBS to a final concentration of 1 mg/mL, filtered and stored at 4°C before performing the cFAE protocol.

### Generation of bsAbs by controlled Fab-arm exchange (cFAE)

BsAbs were generated using a controlled Fab-arm exchange protocol, as reported previously ^34^. Briefly, a Quickchange site-directed mutagenesis kit (New England Biolabs) was used to introduce the F405L and K409R mutations in the HC plasmids of the IgGs to make the bsAbs. A freshly prepared 2-mercaptoethylamine (2-MEA; Sigma) solution (750 mM, pH 7.4) was then mixed with equimolar amounts of IgG1 bearing either the F405L or K409R mutations (1 mg/mL) on a rotating laboratory mixer at room temperature. The mixture was incubated at 31°C for 5 h, and then buffer exchanged to PBS using 100 kDa Vivaspin6columns (Sartorius) to remove the 2-MEA. The mixes were stored at 4°C overnight to allow for reoxidation of the disulfide bonds between the antibody chains.

### Biolayer interferometry (BLI)

To measure the strength of binding of mAbs and bAbs, protein A biosensors (Sartorius) were first loaded with 10 μg/mL of mAb/bsAb in the running buffer until a binding threshold of 1 nm was reached. Following a 60 s wash in running buffer, the biosensors were dipped in a well with SARS-CoV-2 WT, Omicron BA.2, Omicron BA.4/5, SARS-CoV, SHC014, WIV1, Pangolin GX 2017, Rf1 or Khosta-2 S protein (38 μg/mL) for 300 s to measure association, followed by immersion in a well with running buffer for 300 s to measure dissociation. The same method was used to measure the binding of COVA2-02 against soluble RBD and NTD proteins. To investigate the epitope of COVA2-02, we performed a BLI competition assay with the following steps: first (“capture”) mAb (10 μg/mL) was loaded to protein A biosensors for 200 s, followed by a 60 s baseline step in running buffer, first association with 6 μg/mL SARS-CoV-2 WT RBD protein (300 s) and an immediate subsequent association of second (“competitor”) mAb (10 μg/mL) for 200 s, to determine residual binding of mAb to RBD. To test ACE2 competition, we used Ni-NTA sensors (Sartorius), which were loaded with 10 μg/mL of His-tagged SARS-CoV-2 WT RBD for 300 s, followed by a first association with 10 μg/mL of Ab for 300 s and a second association of 10 μg/mL of ACE2 receptor for 300 s. For the preparation of solutions for all BLI experiments and mock measurements, a running buffer containing 0.02% Tween and 0.1% bovine serum albumin in PBS was used. Sensors were regenerated between measurements with a solution of 10 mM glycine in PBS. All BLI experiments were performed using an Octet K2 instrument (ForteBio).

### Pseudovirus production and neutralization assays

All pseudoviruses used in this study were produced by co-transfecting HEK293T cells with the corresponding plasmid expressing the appropriate S with the pHIV-1NL43 ΔEnv-NanoLuc reporter virus plasmid. Cell supernatants containing the pseudoviruses were harvested 48 h post-transfection, centrifuged at 500 x g for 5 min and filtered through a 0.22 µm PVDF syringe filter. Pseudoviruses were then stored at -80°C until further use. To perform pseudovirus neutralization experiments, HEK293T/ACE2 cells were seeded at day 0 with a density of 20.000 cells/well in 100 µL of medium in poly-L-lysine-coated 96-well plates. At day 1, mAbs and bsAbs were serially diluted in DMEM medium, supplemented with 10% fetal calf serum, a mixture of penicillin/streptomycin (100 U/mL and 100 μg/mL, respectively) and 1x glutamax, mixed with the pseudovirus and incubated for 1 h at 37°C. The mixes containing the pseudovirus and the antibodies were then added to the HEK293T/ACE2 cells, seeded the day before. After 48 h, the pseudovirus-antibody combination was removed, cells were lysed and transferred to half-area 96-wells white microplates (Greiner Bio-One). Luciferase activity of cell lysate was measured using the Nano-Glo Luciferase Assay System (Promega) with a Glomax plate reader (Turner BioSystems).

### Negative stain-electron microscopy sample preparation, data collection, and processing

SARS-CoV-2-6P S protein was incubated with either a 0.5-fold M or 3-fold M excess of COVA2-02 for 1 h at room temperature. Complexes were diluted to 0.02 mg/mL in 1x TBS pH 7.4 and deposited on glow discharged carbon-coated copper mesh grids. The grids were stained for 90 sec with 2% uranyl formate and imaged on a Tecnai T12 Spirit at 120 KeV and a 4K x 4K Eagle CCD camera. The Leginon software ^36^ was used to automate data collection and Appion ^37^ was used to store the images. Particles were picked with DogPicker ^38^ and data were processed in RELION 3.0 ^38,39^ for 2D classification.

### Statistical analysis and visualization

Data visualization and statistical analysis were performed in GraphPad Prism 8.3.0. Structural representation of the SARS-CoV-2 WT S and RBD in combination with the ACE2 receptor and mAbs was made by using the UCSF ChimeraX tool ^40^.

## Results

For bsAbs generation, we selected several broad RBD-specific mAbs, with a focus on candidates categorized into class 3 and class 4 ^35^.

From Class 3, we included mAbs S309, Ly-CoV1404, and CV38-142. Class 3 mAbs primarily target the S309 N343 proteoglycan site on the RBD, as previously described ^12,35,41^. S309, originally obtained from an individual infected with SARS-CoV in 2003, serves as the precursor of the commercially known sotrovimab and has maintained activity against SARS-CoV-2 and many of its variants ^42,43^. Ly-CoV1404 (bebtelovimab) is a potent mAb from the same class and exhibits some degree of activity against Omicron BA.4/5 variants, but lacks efficacy against more recent variants like Omicron BQ.1.1 and XBB.1 ^44–47^. CV38-142 was initially identified as a potent class 3 mAb, but was outperformed by broader mAbs with the emergence of recent SARS-CoV-2 variants ^48,49^.

From Class 4, we selected mAbs CR3022, COVA1-16, and 10-40 ^4,50,51^. Class 4 encompasses mAbs targeting the CR3022 site. The benchmark mAb CR3022 was isolated during the SARS-CoV outbreak in 2003. COVA1-16 and 10-40 are well-described broad mAbs whose footprints overlap significantly, as evidenced by their interaction with the RBD (Figure 1).

**Figure 1.**
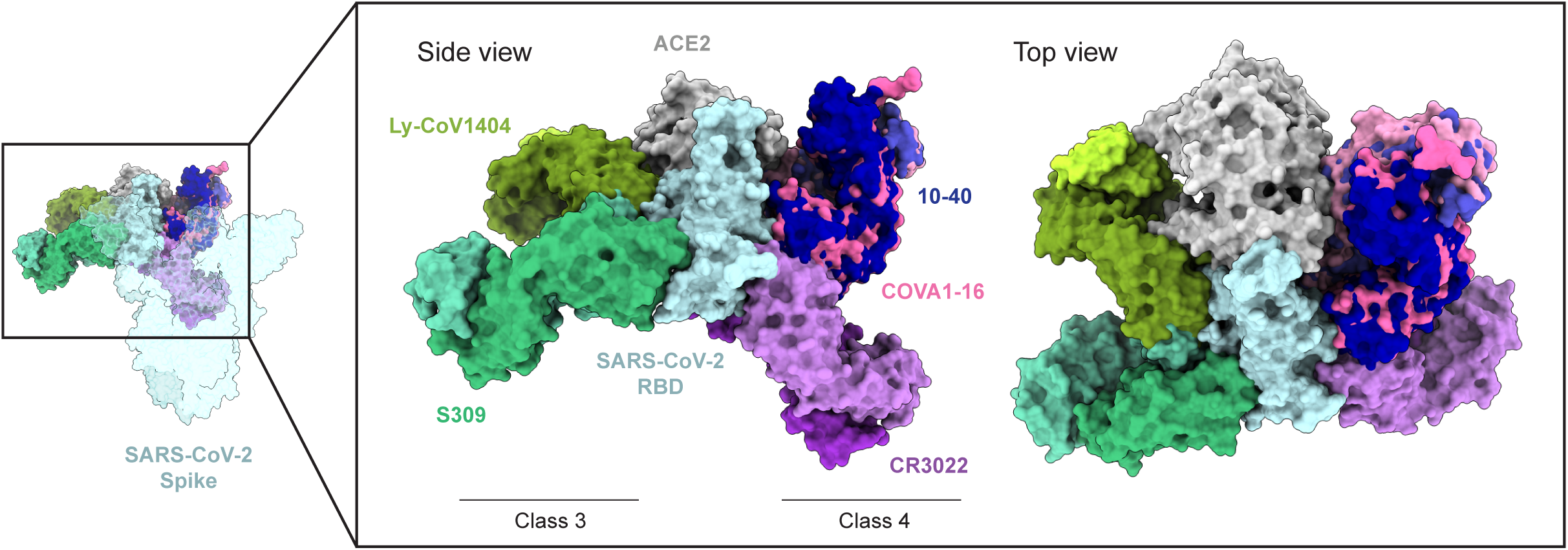
Structural representation of the SARS-CoV-2 RBD with broadly neutralizing mAbs. Depicted is the SARS-CoV-2 S protein (PDB: 7BNN), with focus on one of the three RBD subunits in the “up” conformation, in complex with the Fab regions of mAbs S309 (PDB: 7R6X), Ly-CoV1404 (PDB: 7MMO), CR3022 (PDB: 6YLA), 10-40 (PDB: 7SD5), COVA1-16 (PDB: 7LMW) and the human ACE2 receptor (PDB: 7DF4). A darker color indicates the heavy chain, and a lighter color represents the light chain variable domains.

Additionally, we included mAb COVA2-02, isolated from the COSCA cohort ^4^. This mAb was obtained early in the pandemic and is characterized by its ability to cross-neutralize SARS-CoV and SARS-CoV-2 variants. COVA2-02, when part of a bispecific construct, broadens the neutralization activity of more restricted but potent SARS-CoV-2 mAbs ^4,18^. It targets the RBD, as confirmed by biolayer interferometry (BLI) (Supplementary Figure 1), but its exact binding epitope is unknown.

To produce bsAbs, we combined mAbs with non-overlapping epitopes. Each bsAb contains one arm specific for the class 3 site, whereas the other arm targets the class 4 site. Alternatively, we combined COVA2-02 with either specificities. The binding of all bsAbs and corresponding mAbs was screened by BLI against recombinant sarbecovirus S proteins of SARS-CoV, SHC014 and WIV1 from Clade 1a, SARS-CoV-2 WT, Omicron BA.2, Omicron BA.4/5 and Pangolin GX 2017 from Clade 1b, Rf1 from Clade 2, and Khosta-2 from Clade 3 ^10^ (Supplementary Figure 2). Overall, we observed broad binding for bsAbs containing arms originating from mAbs which also individually bind to S proteins of multiple viruses, such as S309 and COVA1-16. All bsAb combinations including COVA2-02 appeared to have the strongest and broadest binding profile against not only SARS-CoV-2 variants, but also other sarbecoviruses. This can be explained by COVA2-02 recognizing all S proteins included, from weak binding against the Omicron BA.2 S protein to stronger reactivity towards the Rf1 S protein (Figure 2, Supplementary Figure 2). Similarly, the class 4 mAb 10-40, one of the most cross-reactive mAbs described ^4^, was able to bind S proteins from different clades with a comparable or slightly lower affinity than COVA2-02, thus resulting in strong binding of combinations including this mAb.

**Figure 2.**
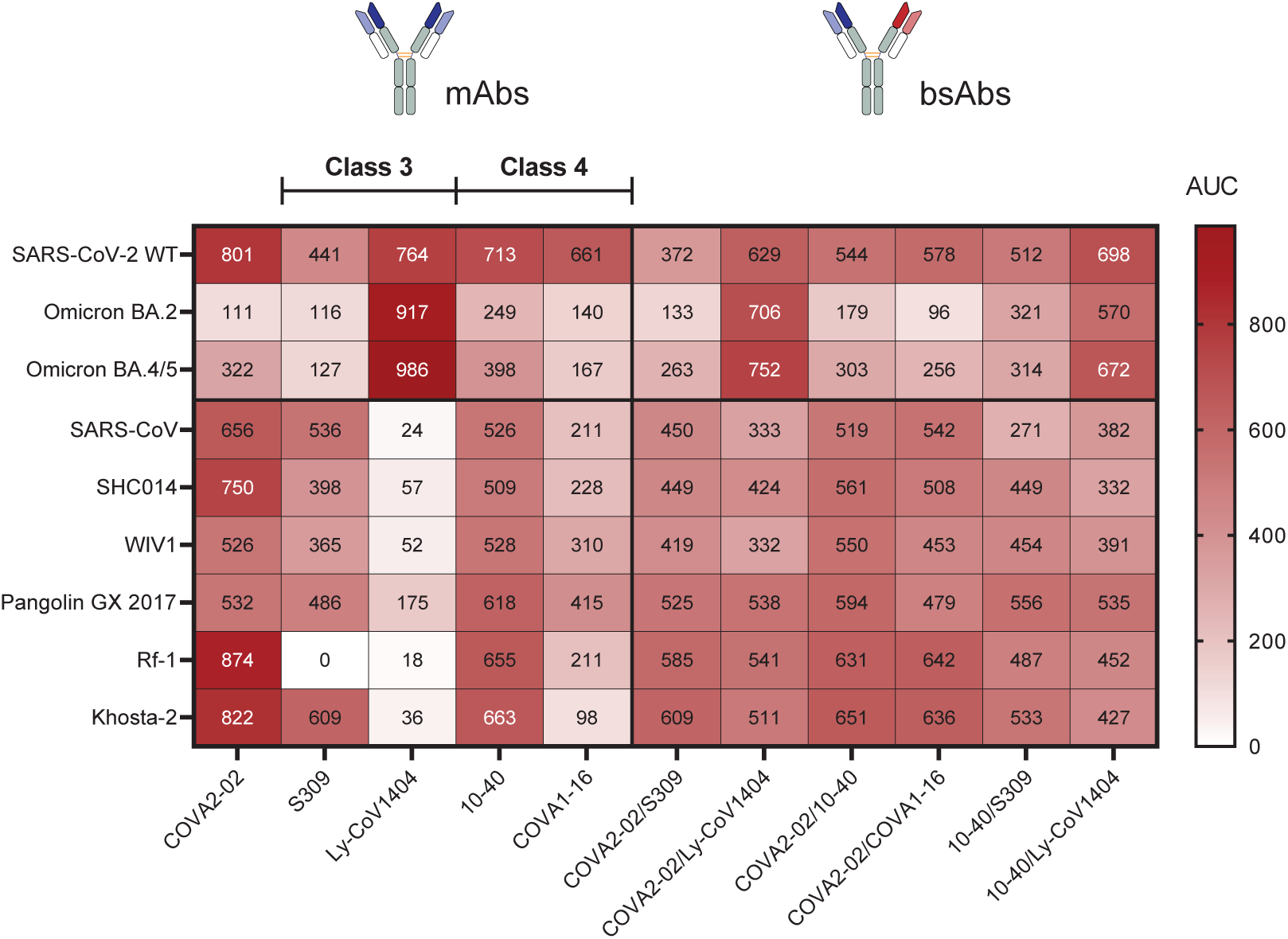
Binding of a subset of bsAbs and corresponding mAbs to several sarbecovirus S proteins. Heatmap showing area under the curve (AUC) values of selected bsAbs including COVA2-02 and other bsAb combinations from class 3 and 4, together with corresponding mAbs, as measured by BLI. S of different SARS-CoV-2 variants (WT, Omicron BA.2 and Omicron BA.4/5), as well as multiple sarbecovirus clades (SARS-CoV, SHC014, WIV1, Pangolin GX 2017, Rf1 and Khosta-2) were included in the assay (distracted from Supplementary Figure 2).

On the contrary, bsAb combinations containing Fab arms of CR3022 and CV38-142 showed narrower specificities and reduced binding. This was also observed for bsAbs containing mAbs isolated from a Gamma-infected individual; COVA309-22, COVA309-35 and COVA309-38 ^20^. Binding of Ly-CoV1404 was restricted to Clade 1b viruses, while bsAbs combining Ly-CoV1404 with 10-40 or COVA2-02 could also bind S proteins of sarbecoviruses from other clades (Figure 2). These results show that binding of non-reactive mAbs can be recovered by combining them with broad binders in a bsAb format.

Next, we examined the pseudovirus neutralization potency of the bsAbs and corresponding mAbs against SARS-CoV-2 WT D614G and the recent variants Omicron BA.4/5, BQ.1.1 and XBB.1, as well as the sarbecoviruses SARS-CoV, WIV1, Pangolin GD 2019 and one merbecovirus, NeoCoV. We first screened all mAbs and bsAb combinations by testing three different concentrations of antibody to determine the neutralization activity (Figure 3A, Supplementary Figure 3A-B). Overall, we observed a trend of increased potency when mAbs are combined into bsAbs. The bsAb neutralization was generally higher, and significantly improved in the case of Pangolin GD 2019 (Mann-Whitney U test, p=0.038). Combining highly specific and narrow mAbs with broader mAbs improved neutralization activity, as we observed for bsAb combinations of narrow mAb Ly-CoV1404 with either broad mAbs 10-40 or COVA2-02. Ly-CoV1404/10-40 and Ly-CoV1404/COVA2-02 neutralized all viruses except NeoCoV with IC_50_ values ranging from 0.002 to 9.5 μg/ml, and from 0.001 to 13.6 μg/ml, respectively, recovering the neutralizing activity of Ly-CoV1404 against Omicron XBB.1, SARS-CoV and WIV1 (Figure 3B, Supplementary Figure 4).

**Figure 3.**
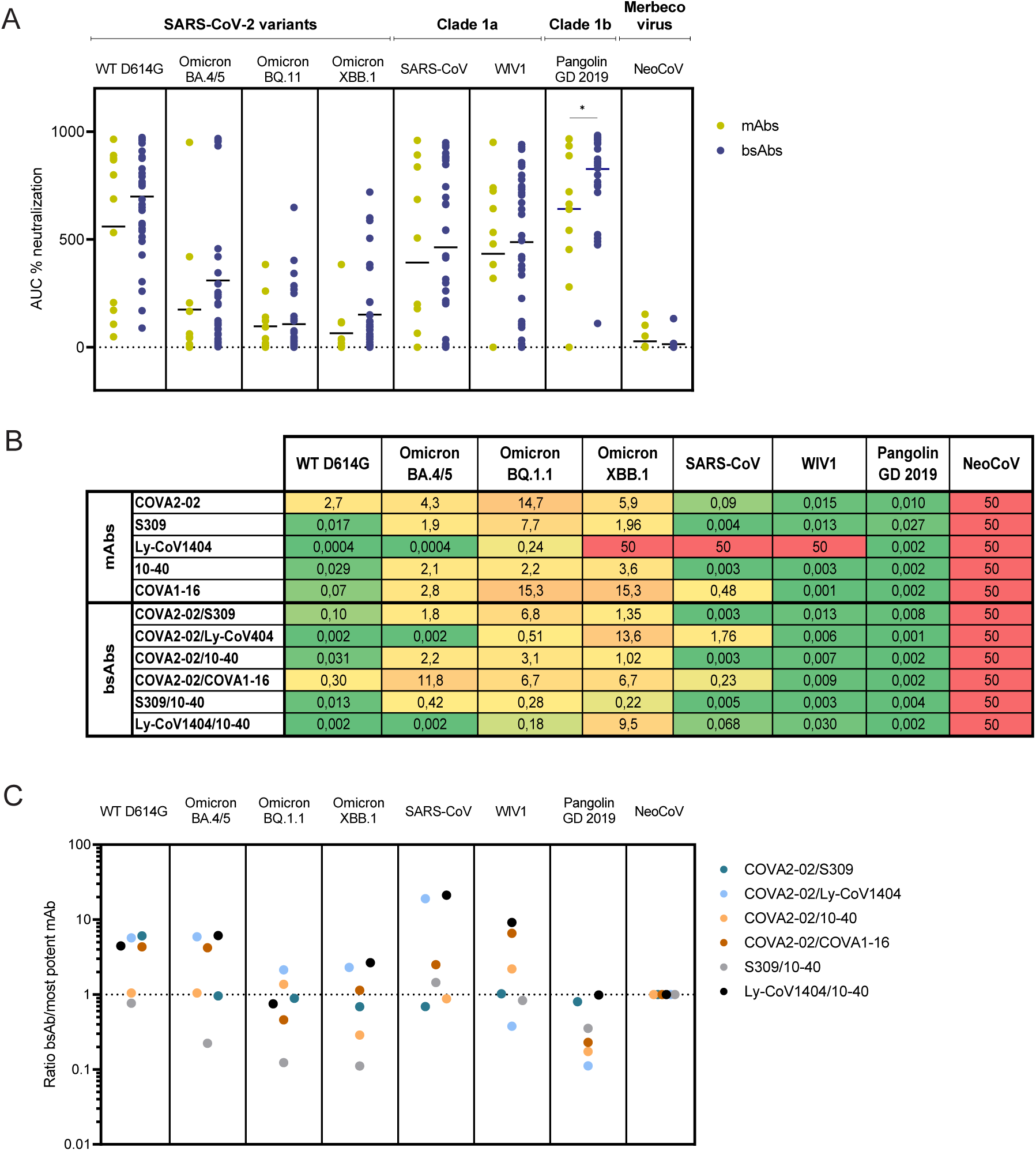
BsAbs neutralization of SARS-CoV-2, its variants and other sarbecoviruses. **A.** Dot graph showing the neutralization for mAbs and bsAbs for the three different concentrations measured in duplo (related to Supplementary Figure 3A-B), and indicated by the % AUC. The mean of all values for a certain group is indicated with a horizontal stripe. Groups were compared using the Mann-Whitney U test, *=p<0.05. Where not indicated, the difference was not statistically significant. **B.** Table showing IC_50_ values of bsAbs including COVA2-02 and corresponding mAbs, and bsAb combinations of class 3 and 4 mAbs, as measured in triplicates by pseudovirus neutralization assay. Each IC_50_ value represents the mean of at least three replicates per mAb/bsAb. **C.** Ratio of IC_50_ values between the bsAbs and the most potent of the two parental mAbs. Values below 1 indicate a significant neutralization improvement of the bsAb combination compared to the mAbs.

When combining broad class 3 and 4 mAbs S309 and 10-40 into a bsAb, we see increased neutralization of all tested SARS-CoV-2 variants compared to both parental mAbs (IC_50_ of 0.42 μg/ml against Omicron BA.4/5, 0.28 μg/ml against Omicron BQ.1.1, and 0.22 μg/ml against Omicron XBB.1, compared to S309 IC_50_s of 1.9, 7.7 and 1.96 μg/ml and 10-40 IC_50_s of 2.1, 2.2 and 3.6 μg/ml, respectively for the three viruses). S309/10-40 bsAb also potently neutralized all tested non-SARS-CoV-2 sarbecoviruses, with IC_50_ values of 0.005, 0.003 and 0.004 μg/ml against SARS-CoV, WIV1 and Pangolin GD 2019, respectively (Figure 3B, Supplementary Figure 4). By comparing our selected bsAb candidates with the more potent of its two parental mAbs (Figure 3C), we conclude that it is possible to increase the individual neutralization activity and achieve cooperative effects when combining potent and broad mAbs into bsAbs.

Interestingly, by replacing either the S309 or 10-40 Fab arm of the S309/10-40 bsAb with a COVA2-02 Fab, the broad neutralizing profile was maintained. COVA2-02/S309 showed an average potency of the two parental mAbs against SARS-CoV-2 WT and retained potency of the more potent mAb against SARS-CoV-2 variants (Figure 3B-C). Importantly, it shows increased activity against non-SARS-CoV-2 sarbecoviruses, such as SARS-CoV (IC_50_ of 0.003 μg/ml), Pangolin GD 2019 (IC_50_ of 0.008 μg/ml), and WIV1 (IC_50_ of 0.013 μg/ml) (Figure 3B, Supplementary Figure 4). COVA2-02/10-40 could potently neutralize most tested viruses, with the highest potency observed against Pangolin GD 2019 (IC_50_ of 0.002 μg/ml). This mAb also shows improvement over both parental antibodies in neutralizing Omicron XBB.1 (Figure 3C). We have previously shown ^18^ that COVA2-02 needs bivalent binding for its activity, as it significantly loses efficacy and neutralization potency when used as functional Fab. However, when we combined COVA2-02 Fab with an active Fab-arm from another mAb in a bsAb format, its affinity to SARS-CoV-2 S protein was restored. We here again demonstrate that when COVA2-02 is included in a bsAb, it can benefit through avidity and synergy to provide modes of binding not available to either one of the two components of the bispecific construct individually.

In light of these findings, we next performed competition BLI experiments to better elucidate the target epitope of COVA2-02 (Figure 4A). When used as capture mAb, COVA2-02 showed no competition with competing mAbs S309, Ly-CoV1404, 10-40, CR3022 or COVA1-16, indicating that it is likely to target a different RBD epitope (Figure 4B). In comparison, when S309 and 10-40 were used as capture mAbs, we observed blocked or reduced consequent binding of Ly-CoV1404, CR3022 and COVA1-16, while COVA2-02 retained its RBD binding capacity (Figure 4B). Next, we performed a BLI experiment to determine whether COVA2-02 competes with the ACE2 receptor for the binding to WT RBD (Figure 4C). COVA2-02, S309 and CR3022 did not compete with ACE2 for binding to the RBD. On the other hand, in line with other studies ^44,50,52^ , Ly-CoV1404, 10-40 and COVA1-16 blocked the RBD binding of the receptor (Figure 4D). These mAbs target epitopes outside the receptor binding site, but their binding causes steric hindrance, making them compete with ACE2. COVA1-18 and COVA2-15, mAbs directly targeting the ACE2 receptor binding site ^4^, also completely blocked ACE2 binding. Overall, these data indicate that COVA2-02 is likely to target a distinct epitope on the lower part of the RBD, which does not overlap with the ACE2 receptor binding site, is not shared by other broad class 3 and 4 mAbs and should therefore be considered when combining mAbs into multispecific antibody formats as a strategy to improve their breadth.

**Figure 4.**
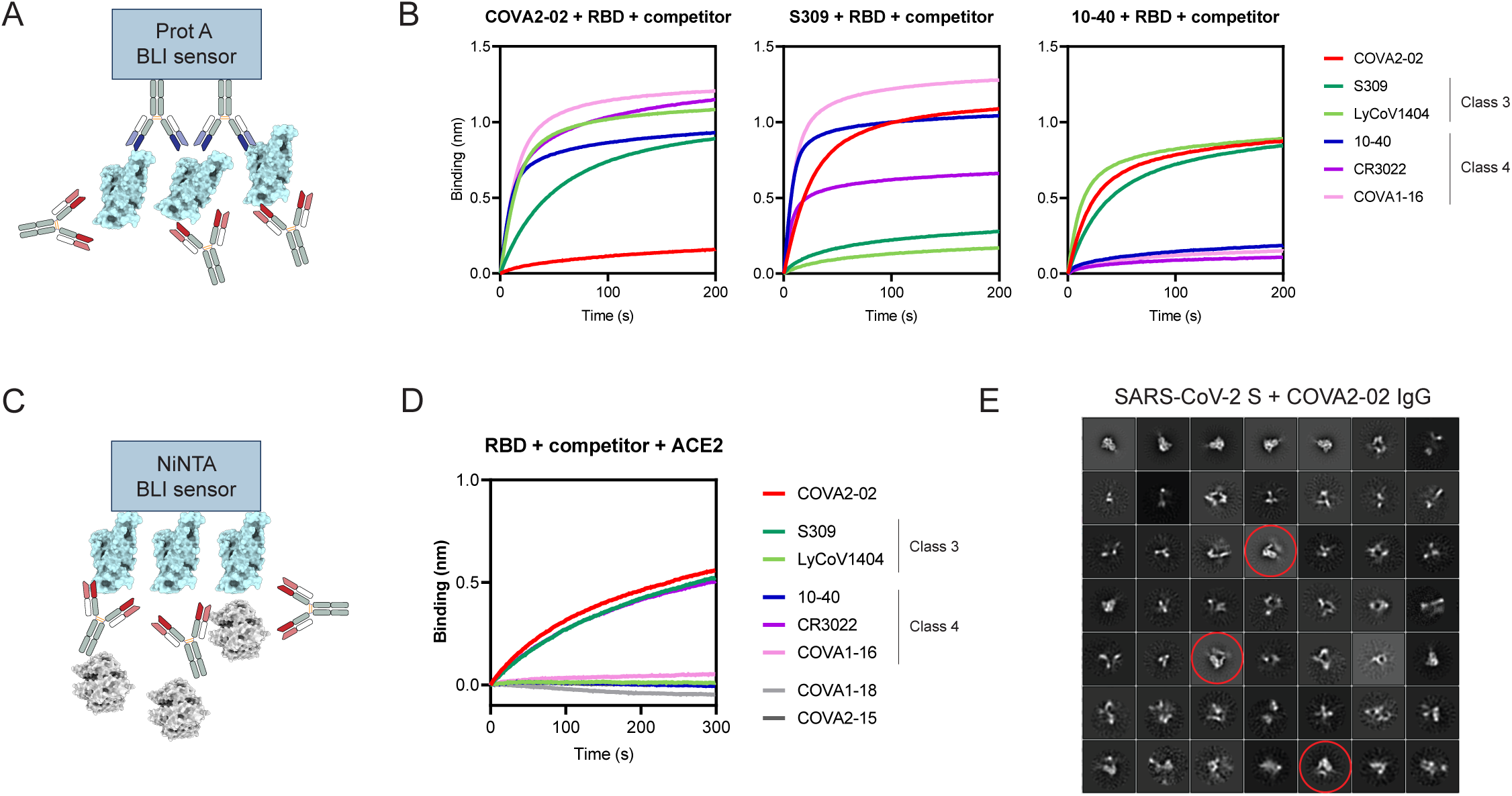
Study of COVA2-02 target epitope. **A.** Schematic setup of the BLI competition experiment with the SARS-CoV-2 WT RBD and known class 3 and 4 mAbs. **B.** BLI competition curves. S309 and 10-40 were used as representative capture mAbs of class 3 and 4, respectively. The curves represent the second association-residual binding of the competitor mAb to the RBD. **C.** Setup of the BLI competition experiment of known class 3 and 4 mAbs and COVA2-02 with the human ACE2 receptor for SARS-CoV-2 WT RBD binding. **D.** BLI competition curves. SARS-CoV-2 WT RBD was loaded onto the biosensor, followed by class 3 and 4 mAbs, or COVA2-02. COVA1-18 and COVA2-15, RBS-targeting mAbs were used as control. The curves represent the second association-residual binding of the human ACE2 receptor to the RBD. **E.** Representative 2D class averages from NS-EM analysis of COVA2-02 IgG bound to SARS-CoV-2 S. Trimer degradation caused by COVA2-02 binding is highlighted with red circles. Due to inherent flexibility and heterogeneity, particles did not converge to a stable 3D class.

Finally, we used negative stain electron microscopy (NS-EM) to better understand the mechanism of action of COVA2-02 IgG targeting the SARS-CoV-2 S protein (Figure 4E). To increase our chances of visualizing the interaction, we used the stabilized SARS-CoV-2-6P S construct. Interestingly, we observed S dimers following COVA2-02 IgG binding, indicating that COVA2-02 is likely to destabilize the S, as was previously reported for other RBD-targeting mAbs ^53,54^. Due to the flexible nature of the S dimer in complex with IgG, the particles were unable to converge in 3D. In addition, we did not observe complexes with NS-EM after incubation of COVA2-02 Fab with S protein (Supplementary Figure 5), indicating a loss of affinity when measuring monovalent (Fab) binding compared to bivalent (IgG) binding.

## Discussion

As has been reported before ^55–57^, the discovery of mAbs that define new conserved and/or cryptic RBD epitopes has an important role in diminishing escape of emerging variants, and informing the development of vaccines that should elicit these types of antibodies. Besides targeting conserved epitopes, many mAbs rely on avidity to efficiently bind the antigen and retain their activity against viral variants, as many SARS-CoV-2 targeting mAbs became significantly less efficient when the bivalent IgG was turned into Fab ^35,52,58^. The loss of affinity for COVA2-02 Fab compared to the full IgG has been suggested by our NS-EM experiments and confirmed by our previous findings ^18^, indicating that this mAb is dependent on bivalency for successful antigen binding. Similarly, a loss of binding (around 2 orders of magnitude higher KDs for the Fab compared to full IgG) was observed for S309 and Ly-CoV1404 ^44,52^, showing that these mAbs also require both arms to retain their affinity. It has been described that mAbs targeting the trimeric SARS-CoV-2 S can exploit the high density of this protein on the surface of the virion for inter-S binding, or bivalently bind one or more of the three RBDs at the same time, for intra-S cross-linking ^18,35,58^. Although by creating bispecific constructs the bivalency of each parental mAb is sacrificed, we demonstrate here that by combining individual Fabs into a bsAb, binding can be recovered. BsAbs thus allow not only for variety in target epitopes contained in one molecule, but also a combination of binding mechanisms which can provide beneficial cooperativity.

Most of the bsAb combinations produced in this study had retained or even increased binding strength to S proteins of different sarbecoviruses. Others have reported synergistic neutralization of SARS-CoV-2 variants combining broad class 4 mAbs with more specific class 1 mAbs ^59^, and similarly to this study, class 3 and 4 mAbs ^48,60^. Of note, in a previous study ^48^, a mix of class 3 CV38-142 and class 4 COVA1-16 showed modest synergistic effects. In our study, this bsAb was significantly outperformed by other combinations, such as S309/10-40 and Ly-CoV1404/10-40. BsAbs containing COVA2-02 were also highly efficient and broad neutralizers, sometimes surpassing the potency of both COVA2-02 and the other individual parental mAb, by up to 8-fold (in the case of COVA2-02/Ly-CoV1404 against Pangolin GD 2019). This indicates that COVA2-02 is likely to benefit from a synergistic effect with other SARS-CoV-2-specific mAbs, presumably by conformational changes upon binding of the first Fab, or by stabilization of one of the three RBDs in an open state to make additional epitopes available. We have previously shown this by combining COVA2-02 with COVA2-15, a very potent but narrow class 2 mAb, into a bsAb with improved potency against SARS-CoV-2 and several earlier variants ^18^. Previously we also showed superiority of bsAbs containing COVA2-02 compared to corresponding cocktails, and demonstrated stoichiometries of S protein binding for the bsAb unavailable to mAbs or cocktails ^18^. Overall, our results suggest that COVA2-02 has the potential to cooperate and synergize in neutralization with other mAbs, and could be a good candidate for future bsAb therapeutic formulations. We emphasize the importance of addressing avidity of mAbs as a strategy to overcome the emergence of viral variants and we also corroborate the hypothesis of combining broad mAbs targeting epitopes closer to the base of the RBD into bsAbs.

Another advantage of bsAbs over cocktails of mAbs can be the potentially simplified development process and reduced manufacturing costs. However, for these benefits to be feasible in practice, the production of bsAbs needs to be simple, high-throughput and flexible in terms of quick adaptation of selected mAb specificities to the current variant landscape. The generic cFAE method has been proven to be a feasible technology, with several bispecific candidates in clinical development and one mAb, amivantamab, approved by the Food and Drug Administration to treat non-small cell lung cancer ^34,61^. Besides the modular versatility of this platform to easily combine different antibody specificities into one molecule, the method could be further simplified by adjusting protocols so that parental mAbs can be co-transfected and a full cFAE produced antibody can be purified directly from the supernatant ^62^ Alternatively, this method could be expanded by incorporating VHH-molecules at the C-termini of bispecific IgGs, to produce tri- and tetravalent constructs ^63^. By increasing the valency, the potency and breadth could be further improved.

In addition, we have previously shown that bsAbs generated by cFAE retained *in vitro* antibody-dependent cellular phagocytosis and trogocytosis ^18^, and expect the bsAbs described in this study to retain these properties as well. However, particularly mAbs that do not use direct ACE2 blocking as their mechanism of neutralization, often rely on S destabilization, conformational changes and Fc-mediated effector functions to combat viral infection ^42^. Therefore, the presence of antibody-dependent cellular cytotoxicity, phagocytosis and trogocytosis should be determined for candidates proceeding to further *in vitro* and clinical studies.

In conclusion, this study highlights the need for characterizing new cross-neutralizing antibodies and potential synergistic mAb combinations, to create additional prophylactic agents and antibody therapeutics. This is of particular value for combating breakthrough infections and potential new global outbreaks caused by zoonotic spillover of sarbecoviruses to the human population.

## Acknowledgments

We thank Dirk Eggink of the National Institute for Public Health and the Environment (RIVM), Bilthoven, the Netherlands, for providing the Omicron BA.2 and BA.4/5 S protein constructs.

## Author contributions

Conceptualization: D.G., L.R., T.B., K.S. and M.J.vG.

Methodology: D.G., L.R. and K.S.

Investigation: D.G., L.R., M.P., M.B., L.v.d.M. and J.L.T.

Resources: D.G., L.R., M.P., M.B., L.v.d.M., J.L.T., T.B. and M.J.vG.

Data curation and visualization: D.G., L.R., L.v.d.M. and J.L.T.

Supervision: T.B., K.S., J.S., M.J.vG. and R.W.S.

Funding acquisition: J.S., M.J.vG., A.B.W and R.W.S.

Writing-Original draft: D.G., L.R. and T.B.

Writing-Review and editing: all authors.

## Declaration of interest

None of the authors have conflicts of interest related to this research.

## Funding

This work was supported by a Netherlands Organisation for Scientific Research (NWO) Vici Grant (no. 91818627) to R.W.S, by the Fondation Dormeur, Vaduz to R.W.S and M.J.vG., and by a Vidi and Aspasia grant from the NWO (grant numbers 91719372 and 015.015.042) to J.S.

---

**Supplementary Figure 1.**
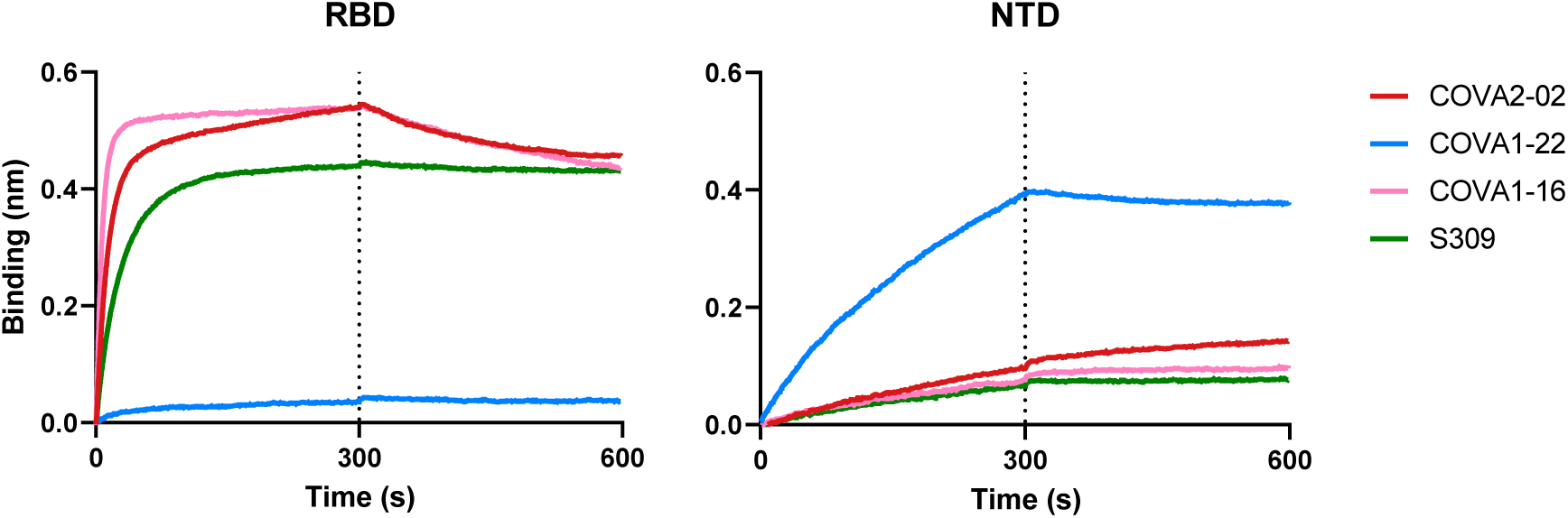
COVA2-02 binding to SARS-CoV-2 RBD. BLI affinity measurements of COVA2-02, control RBD-binding mAbs COVA1-16 and S309, and a NTD-binding mAb COVA1-22 to SARS-CoV-2 RBD (left) or NTD (right).

**Supplementary Figure 2.**
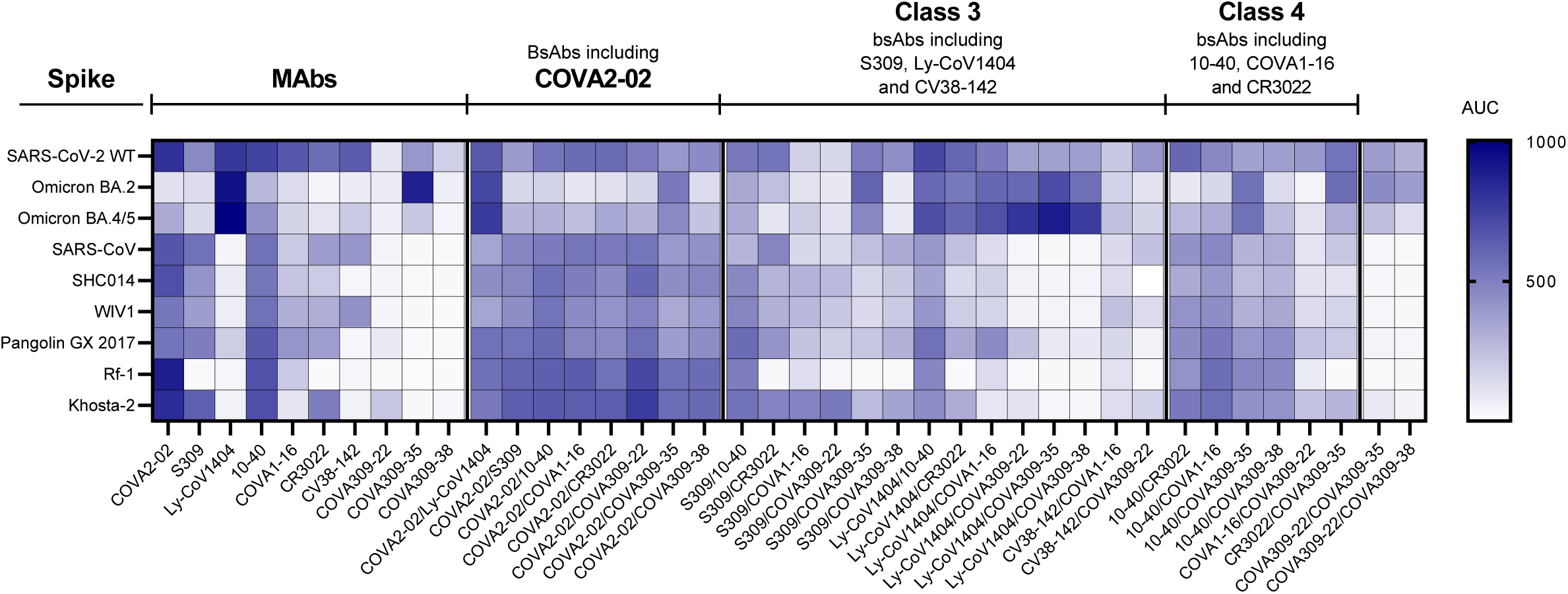
BLI affinity measurements of all the bsAbs and corresponding mAbs against S from different sarbecoviruses. Samples were grouped as follows: mAbs, bsAbs with one COVA2-02 arm, bsAbs including one class 3-specific arm, bsAbs including one class 4-specific arm. The heatmap shows AUC values.

**Supplementary Figure 3.**
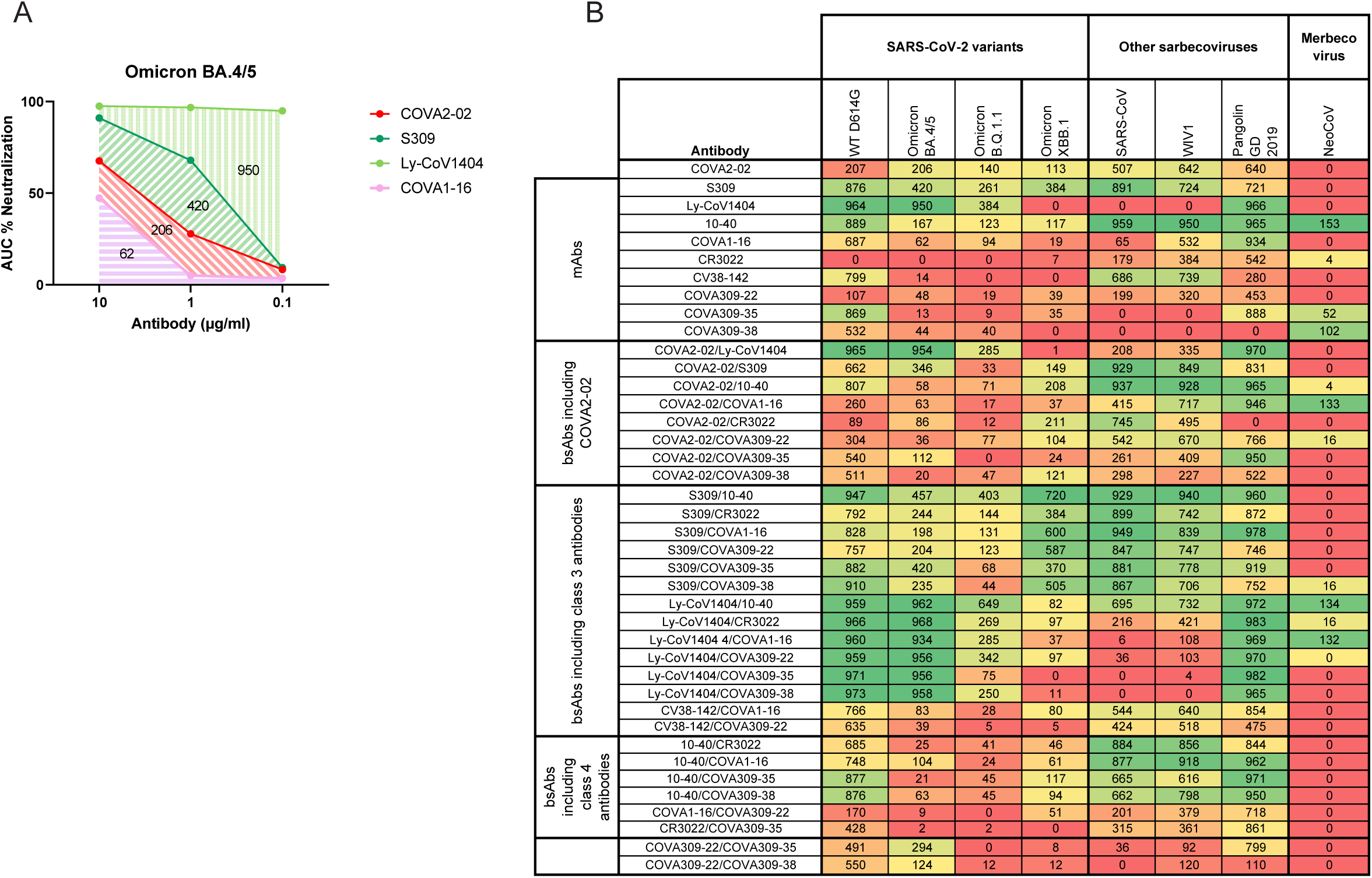
Pseudovirus neutralization measurements of mAbs and bsAbs against sarbecoviruses. **A.** Representative graph showing the percentage of neutralization indicated by the AUC obtained after measurements using three concentrations (10, 1 or 0.1 µg/mL) of mAb or bsAb against Omicron BA.4/5 pseudovirus. AUC values are indicated as numbers on the graph for each antibody. All AUCs were calculated taking into account only values above 0. All values are baseline subtracted and normalized. **B.** Table showing raw values of AUC of percentage neutralization for all measured mAbs and produced bsAbs against 8 different viruses.

**Supplementary Figure 4.**
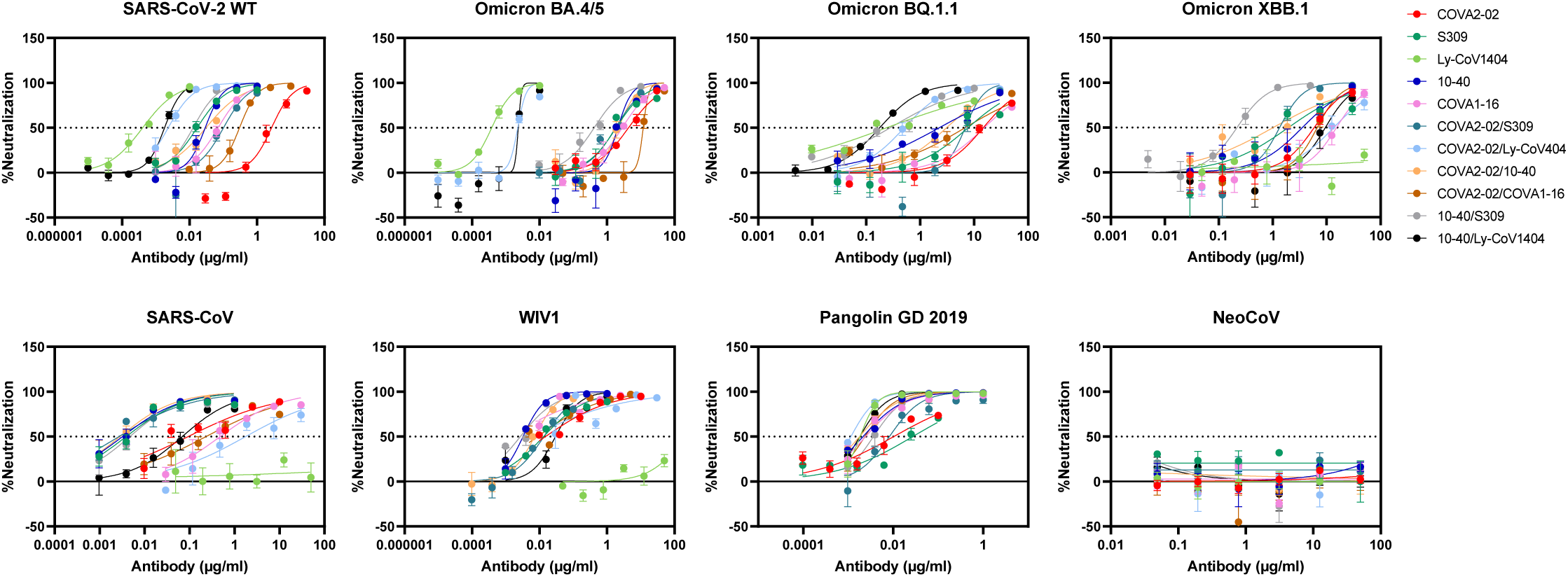
BsAbs and mAbs neutralization of different sarbecoviruses. Curves of bsAbs and corresponding mAbs indicated by the percentage of neutralization of SARS-CoV-2 and its variants (top panel), as well as other sarbecoviruses and one merbecovirus (bottom panel). Dot lines indicate the antibody concentration at which 50% of neutralization is achieved. The assay was performed in triplicates.

**Supplementary Figure 5.**
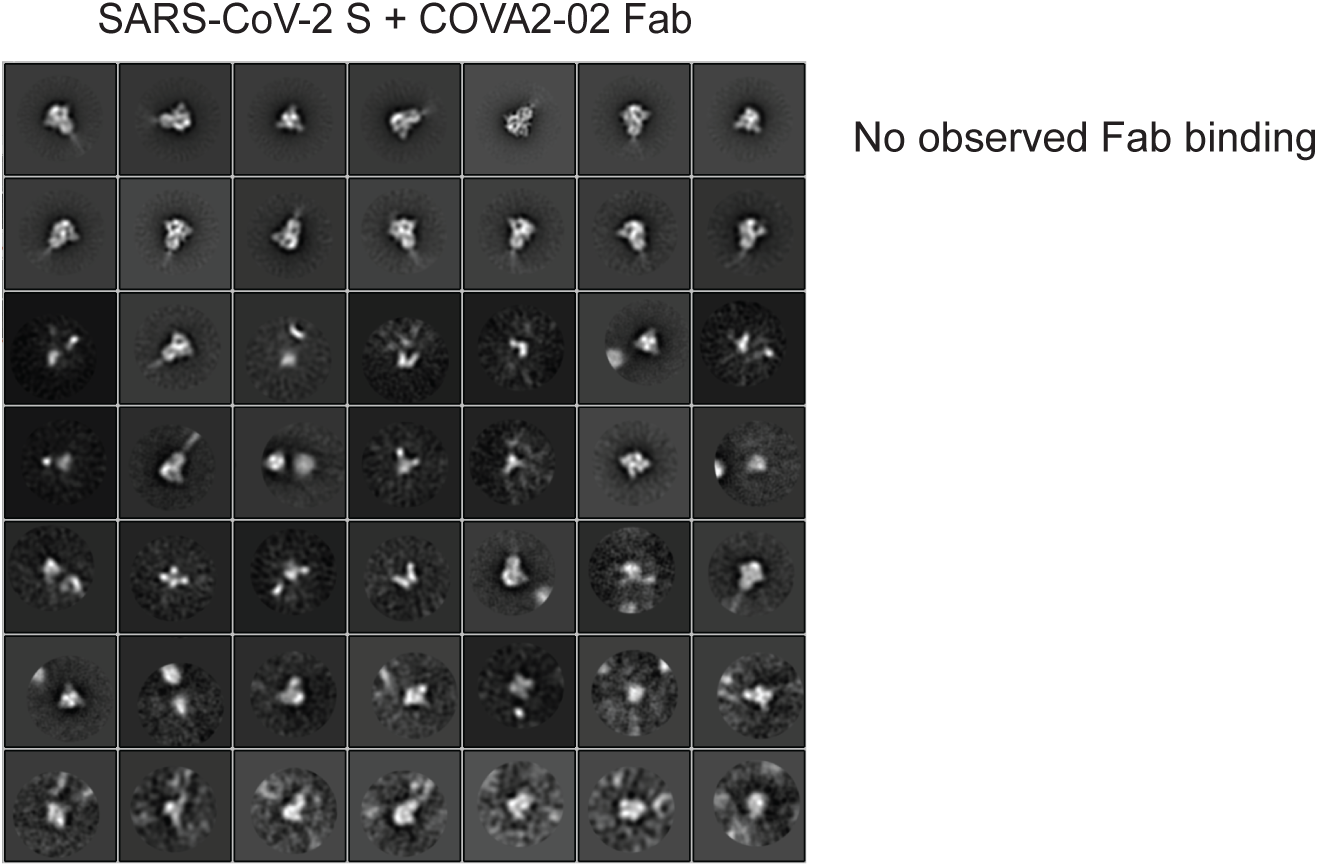
Representative 2D class averages from NS-EM analysis of COVA2-02 Fab. COVA2-02 Fab was incubated with SARS-CoV-2 S, but no complexes were observed.

## Notes

### Competing Interest Statement

The authors have declared no competing interest.

